# Generation and validation of a D1 dopamine receptor Flpo knock-in mouse

**DOI:** 10.1101/2024.04.25.591164

**Authors:** Alexis M. Oppman, William J. Paradee, Nandakumar S. Narayanan, Young-cho Kim

## Abstract

**Background:** Dopamine is a powerful neuromodulator of diverse brain functions, including movement, motivation, reward, and cognition. D1-type dopamine receptors (D1DRs) are the most prevalently expressed dopamine receptors in the brain. Neurons expressing D1DRs are heterogeneous and involve several subpopulations. Studying these neurons has been limited by current animal models, especially when considering their integration with conditional or intersectional genetic tools.

**New method:** To address this limitation, we developed a novel Drd1-P2A-Flpo (Drd1-Flpo) mouse line in which the Flpo gene was knocked in immediately after the Drd1 gene using CRISPR-Cas9. We validated the Drd1-Flpo line by confirming Flp expression and functionality specific to D1DR+ neurons.

Comparison with existing methods: The Drd1-Flpo line is useful resource for studying subpopulation of D1DR+ neurons with intersectional genetic tools.

**Conclusions:** We demonstrated brain-wide GFP expression driven by Drd1-Flpo, suggesting that this mouse line may be useful for comprehensive anatomical and functional studies in many brain regions. The Drd1-Flpo model will advance the study of dopaminergic signaling by providing a new tool for investigating the diverse roles of D1DR+ neurons and their subpopulations in brain disease.

**Significance Statement:** The roles of dopamine in the brain are mediated by dopamine receptors. D1-type dopamine receptors (D1DRs) and D1DR-expressing (D1DR+) neurons play important roles in various brain functions. We generated a Drd1-Flpo mouse line that expresses Flp recombinase in D1DR+ neurons. This novel Drd1-Flpo mouse facilitates investigation of specific roles of D1DR+ neurons in various brain areas including the striatum, frontal cortex, and cerebellum, and it provides an alternative to existing Drd1-Cre mice. In addition, the Drd1-Flpo mouse line provides a tool for intersectional genetic studies, when used with existing transgenic Cre lines. The Drd1-Flpo mouse line can help unravel the specific contributions of D1DR+ neuron subpopulations to brain function and dysfunction.

## INTRODUCTION

Dopamine plays an important role in human physiology by modulating functions such as movement, motivation, reward, and cognition (Klein et al., 2019). Dopamine modulates neuronal signaling primarily via two broad classes of receptors—D1-like and D2-like dopamine receptors. These are G-protein-linked receptors that have an opposite effect on adenylyl cyclase (Neve et al., 2004; Kim et al., 2015). Five identified dopamine receptors, D1DR, D2DR, D3DR, D4DR, and D5DR, are encoded by five distinct genes, Drd1, Drd2, Drd3, Drd4, and Drd5, respectively. D1DRs are the most abundant type of dopamine receptors in the brain, and D1DR-expressing (D1DR+) neurons are involved in motor function and cognitive processes such as working memory, spatial learning, and temporal processing (Clark and White, 1987; Sawaguchi and Goldman-Rakic, 1991; Yang et al., 2018; Kim and Narayanan, 2019; Matzel and Sauce, 2023). Further exploration and characterization of D1DR+ neurons may facilitate new targeted therapies for motor and cognitive impairments of human diseases such as Parkinson’s disease, schizophrenia, and Alzheimer’s disease (Kim et al., 2015; Anastasiades et al., 2019; Lamptey et al., 2022).

D1DR+ neurons are distributed throughout many brain regions, including the frontal cortex, thalamus, and amygdala, and on medium spiny neurons of the dorsal and ventral striatum (Wei et al., 2018). Within these areas, D1DRs are expressed in distinct cell populations such as pyramidal neurons and interneurons (Glausier et al., 2009). Genetically-modified mouse lines facilitate specific studies of D1DR+ neurons (Kravitz et al., 2010; Gangarossa et al., 2012; Han et al., 2017; Kim et al., 2017; Kim and Narayanan, 2019). To further dissect D1DR+ circuits— such as identifying the roles of D1DR+ pyramidal neurons vs. D1DR+ interneurons in the cortex, or to distinguish between D1DR+ vs. D2DR+ neurons in the striatum—more specific tools are needed. Ideally, these tools would enable conditional expression or intersectional genetics allowing D1DR+ neurons to be manipulated with multiple levels of control (Taniguchi et al., 2011; Fenno et al., 2014; Poulin et al., 2018). For instance, targeting D1DR+ neurons in pyramidal neurons would require at least two levels of control to isolate D1DR+ neurons specifically in pyramidal neurons.

We developed and validated a novel transgenic Drd1-Flpo mouse generated using CRISPR-Cas9. To verify Flp expression and functionality, we expressed mCherry using an Flp-conditional viral vector injected in the prefrontal cortex (PFC) of Drd1-Flpo mice. To identify Flp expression brain-wide, we crossed Drd1-Flpo mice with an Flp-conditional enhanced green fluorescent protein (eGFP) reporter mouse line (FRT-eGFP). We then imaged cryo-sectioned brain slices and tissue-cleared brains to quantitatively image whole-brain Drd1-Flpo/FRT-eGFP expression. Our findings advance a novel transgenic tool to specifically manipulate D1DR+ neurons and circuits in combination with conditional and intersectional genetic strategies.

## METHODS

### Animals

All procedures were approved by the University of Iowa Institutional Animal Care and Use Committee (IACUC) in accordance with the established guidelines and regulations (Protocol #0062039). We used a total of 9 mice, including 1 Drd1-Flpo (1 males) and 8 Drd1-Flpo/FRT-eGFP (3 females, 5 males) mice in this study. All mice were bred and housed under a 12-hour light-dark cycle, with ad libitum access to food and water.

### Drd1-Flpo Knock in genetic engineering

We used CRISPR-Cas9 to generate D1-type dopamine receptor Flp recombinase mice (Drd1-Flpo) in collaboration with the Genome Editing Facility at the University of Iowa. Male mice older than 8 weeks were bred with 3–5-week-old super-ovulated females to produce zygotes for pronuclear injection. Female Institute of Cancer Research (ICR) mice (Envigo, Indianapolis, IN; Hsc:ICR(CD-1)) were used as recipients for embryo transfer. Pronuclear-stage embryos were collected using methods described previously (Pinkert, 2014). Embryos were collected in KSOM media (Millipore Sigma, Burlington, MA; MR101D) and washed 3 times to remove cumulous cells. Cas9 ribonucleoprotein (RNP) and double-stranded repair template were injected into the pronuclei of the collected zygotes and incubated in KSOM with amino acids at 37 °C under 5% CO2 until all zygotes were injected. Fifteen to 25 embryos were immediately implanted into the oviducts of pseudo-pregnant ICR females.

Chemically modified CRISPR-Cas9 crRNA and CRISPR-Cas9 tracrRNA were purchased from Integrated DNA Technologies (IDT) (Coralville, IA; Alt-R^®^ CRISPR-Cas9 crRNA; Alt-R^®^ CRISPR-Cas9 tracrRNA; Catalog #1072532). The crRNA and tracrRNA were suspended in T10E0.1 buffer and combined to a 1 ug/ul (∼29.5 uM) final concentration in a 1:2 (ug:ug) ratio. The RNAs were then heated at 98 °C for 2 minutes and allowed to cool slowly to 20 °C in a thermal cycler. The annealed cr:tracrRNAs were aliquoted to single-use tubes and stored at -80 °C . Cr:tracr:Cas9 RNP complexes were made by combining Cas9 protein (IDT; Alt-R^®^ S.p. HiFi Cas9 Nuclease) and cr:tracrRNA in T10E0.1 buffer (final concentrations: 300 ng/ul (∼1.9 uM) Cas9 protein and 200 ng/ul (∼5.9 uM) cr:tracrRNA). The Cas9 protein and annealed RNAs were incubated at 37 °C for 10 minutes. These RNP complexes were combined with single-stranded repair template and incubated an additional 5 minutes at 37 °C. The concentrations in the injection mix were 30 ng/ul (∼0.2 uM) Cas9 protein; 10 ng/ul (∼0.3 uM) cr:tracrRNA; and 10 ng/ul single-stranded repair template. A custom 6120-bp DNA sequence (Drd1_Guide_B, GGTTGAATGCTGTCCGCTGT) containing P2A-Flpo (ATNFSLLKQAGDVEENPGP, SV40 nuclear localization sequence), with a 5’ homology arm (686 bp) and a 3’ homology arm (617 bp), was inserted at the 2901 bp in the Drd1a gene, immediately upstream of the stop codon in exon 2, generating the resulting targeted insertion allele. During Drd1a gene translation, the P2A site within the insertion prompts self-cleavage, resulting in the separation of Drd1a from Flpo (Szymczak et al., 2004). Consequently, Drd1a proceeds with regular migration to the cell membrane, while Flp remains confined within the cytoplasm. We verified mouse genotypes using primers for the Drd1-Flpo recombinase transgene (523 bp) (Drd1a-Flpo-F: 5’-GAC TCT GCC CTA CAA CGA AT-3’; Drd1a-Flpo-R: 5’-GTG ATC ATC CAG CAC AGG T-3’).

### Validating Flp expression

To investigate the functionality of the P2A-Flpo insertion, we stereotaxically injected a Drd1-Flpo animal with AAV1-Ef1a-fDIO-mCherry (Addgene, Watertown, MA; Catalog #114471-AAV1), an adeno-associated virus (AAV) with an Flp-conditional switch that expresses mCherry, driven by the Ef1a promoter. The animal was injected with 1 µL of the AAV virus in the medial PFC (mPFC; AP: +1.8, ML: -0.2, DV: -1.9 and -1.7). Following an incubation period of at least 21 days, the animal was euthanized and perfused.

To examine brain-wide expression of Flp, Drd1-Flpo animals were crossed with a homozygous FRT-eGFP reporter mouse strain (STOCK Gt(ROSA)26Sortm1.2(CAG-eGFP)Fsh/Mmjax; Jackson Laboratory, Bar Harbor, ME), whose endogenous eGFP expression is dependent on FRT-flanked *STOP* cassette excision by Flp. Homozygous FRT-eGFP mouse genotypes were verified using primers for the Drd1-Flpo recombinase transgene.

### Immunohistochemistry

Mice were anesthetized by intraperitoneal injection with 100 mg/kg ketamine (Dechra Veterinary Products, Overland Park, KS) and 100 mg/kg xylazine (AnaSed Injection, Akorn, Inc. Lake Forest, IL). Mice were intracardially perfused for 5 minutes with ice cold 1x DPBS followed by 5 minutes of ice cold 4% paraformaldehyde (PFA; Thermo Scientific, Fair Lawn, NJ). Following perfusion, brains were immediately dissected and incubated overnight in 4% PFA at 4 °C. Brains were transferred to 30% sucrose in 1x DPBS for 48 hours and frozen in O.C.T. embedding media (Scigen FisherHealthCare, Houston, TX) at -80 °C. Brains were sectioned coronally at 40µm on a cryostat (Leica Biosystems, Deer Park, IL).

For immunohistochemistry, free floating sections were blocked for 1 hour in 5% normal goat serum in 0.3% PBST (Triton-X 100 in PBS) at 4 °C. The diluted primary antibody was applied and incubated overnight at 4 °C. A rat anti-D1DR monoclonal antibody was used for D1DR-receptor staining (1:500; Sigma-Aldrich, St Louis, MO; Catalog #D2944). To enhance the GFP signals, a chicken anti-GFP antibody (1:500; Abcam, Cambridge, UK; Catalog #AB13970) was used. Following overnight incubation, sections were washed with 0.3% PBST, and diluted secondary antibodies were applied at 4 °C for approximately 3 hours using Alexa Fluor 488 goat anti-rabbit (1:1000; Invitrogen, Waltham, MA; Catalog #A11008); Alexa Fluor Plus 555 goat anti-rat (1:1000; Invitrogen; Catalog #A48263); and Alexa Fluor 488 goat anti-Chicken (1:1000; Invitrogen; Catalog #A11039). Sections were incubated either in DAPI (Life Technologies, Carlsbad, CA) in PBS for 5 minutes before mounting or directly mounted in anti-fade mounting media containing DAPI. Sections were stored at 4 °C until imaging with an Olympus fluorescent microscope (VS120; Olympus Life Science, Waltham, MA).

### Microscopy and colocalization quantification

Using an Olympus VS120 slide scanning microscope (Evident, Tokyo, Japan), images were acquired with 20x objectives with 21 (1.8-um step) z-spaced images. The z-stacked images were 3D-deconvoluted with the *deconvlucy* algorithm (Li et al., 2018), and then z-projected images were generated with an extended depth of field algorithm (Pertuz et al., 2013). We used custom MATLAB (Mathworks, Natick, MA) codes for colocalization analysis of anti-GFP and anti-D1DR immunostaining. GFP channel images were filtered with a 2D gaussian kernel before being converted to binary images to identify GFP-expressing cell bodies. The clean shape of cell bodies was identified using the Chan-Vese active contour (Chan and Vese, 2001). The anti-D1DR channel was converted to binary images with the adaptive image binary function, *imbinarize*, in MATLAB. Two binary images were processed with the *regionprops* function to identify common pixels.

### Whole-brain tissue clearing and imaging

Tissue clearing methods were used to quantify whole-brain D1DR and GFP expression in two Drd1-Flpo/FRT-eGFP animals. Mice were perfused as previously described, and subsequent processing, including SHIELD fixation preprocessing and delipidation (for clearing with stochastic electrotransport), tissue clearing, immunolabeling, and microscopy, was performed at LifeCanvas Technologies (Cambridge, MA) (Murray et al., 2015; Park et al., 2019; Yun et al., 2019). Immunolabeling was performed using goat anti-GFP (8 ug/brain; EnCor, Gainesville Florida; #GPCA-GFP); rat anti-D1DR (8 ug/brain sample; Sigma-Aldrich; D2944), and rabbit anti-NeuN (6 ug/brain sample; Cell Signaling Technology, Danvers, MA; #24307) antibodies. Imaging at 3.6x magnification with 1.8-um xy steps and 4-um z-steps was performed with a SmartSPIM light sheet microscope (Bruker, Billerica, MA), and images were registered to the Allen Brain Atlas (Allen Institute, Seattle, WA). A NeuN channel for each brain was registered to an average NeuN atlas. Registration was performed using successive rigid, affine, and b-spline warping algorithms (SimpleElastix: https://simpleelastix.github.io/). Cell detection for the GFP channel was performed, and Life Canvas Technologies provided all raw image data and analysis outputs, including cell count CSV files, and heat maps of the data. The raw TIF files were converted to a 3D-mapped matrix and resliced coronally or sagittally, then encoded to MP4 video files in MATLAB. Using the registered images, intensity of the 488-nm and 561-nm channels was linearly fitted in 838 brain areas (average of both hemispheres, 2 mice) to examine correlation between D1DR antibodies and Flp-dependent GFP expression. All code and raw data are available at https://narayanan.lab.uiowa.edu.

## RESULTS

### Generation of Drd1-Flpo transgenic mice

Our goal was to develop a mouse line that would enable cell-type-specific manipulation of D1DR+ neurons. We generated a Flp recombinase-expressing mouse line by inserting a P2A-Flpo sequence at the endogenous Drd1a genomic locus with CRISPR-Cas9 knock-in techniques (Fig 1A). The P2A peptide sequence allows Drd1 and Flpo to be transcribed as a single mRNA. The P2A peptide sequence self-cleaves upon translation, dissociating the D1DR transmembrane receptor from Flpo recombinase. This strategy allows expression of Flp in the cytoplasm of all cells that express D1DRs.

**Figure 1:**
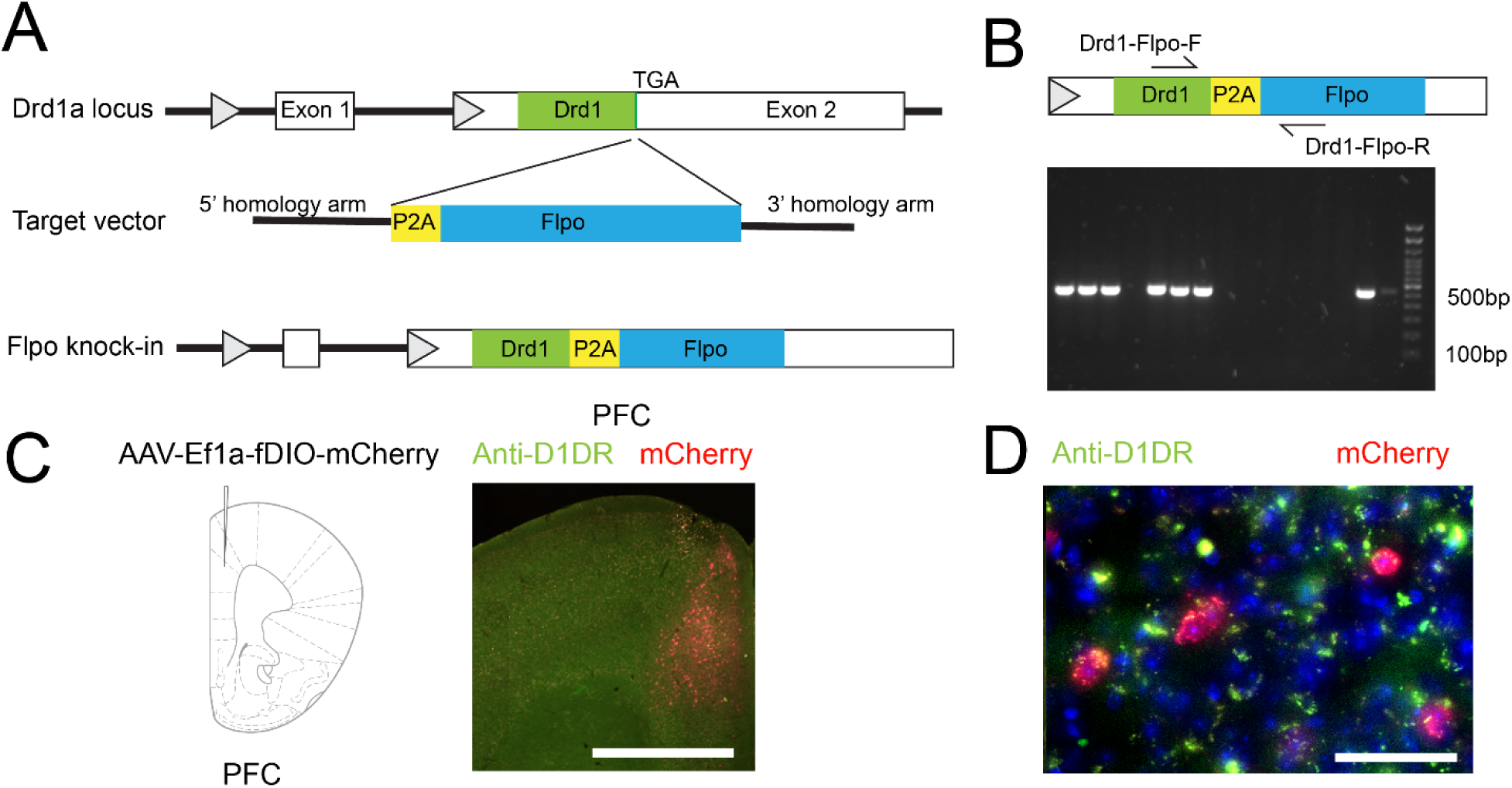
Schematic of Drd1-Flpo transgene creation. (A) Targeting the Drd1a genomic locus to generate Drd1-Flpo knock-in mice. (B) PCR genotyping of Drd1-Flpo mice. The 523-bp band is amplified with primers flanking the Drd1 and Flpo gene. (C) Illustration of injection with AAV-fDIO-mCherry into the rodent prefrontal cortex. Expression of mCherry in the dorsal prelimbic and anterior cingulate cortex of Drd1-Flpo mice (scale bar = 1000 um). D) Cortical colocalization of mCherry (red) and anti-D1DR antibody staining (green) (scale bar = 50 um).

We confirmed the insertion of the targeted sequence using PCR (Fig 1B). We used viral methods to confirm Flpo recombinase expression in D1DR neurons by injecting AAV containing Flp-dependent mCherry (fDIO-mCherry, AAV1-Ef1a-fDIO-mCherry) into the mPFC of Drd1-Flpo mice (prelimbic/anterior cingulate; Fig 1C). Twenty-one days after viral injection, mice were perfused, and brains were processed for immunohistochemistry. We found strong expression of prefrontal mCherry around the viral injection sites (Fig 1C). We co-stained sections with antibodies against D1DRs (anti-D1DR) and using high-power microscopy, found co-expression of mCherry and anti-D1DR (Fig 1D). These data suggest that mCherry was conditionally expressed in Drd1-Flpo mice.

### Drd1-Flpo mice express Flp in D1DR+ neurons

Next, we examined Flp expression in targeted brain areas. We bred Drd1-Flpo mice with FRT-eGFP; Gt(ROSA)26Sor reporter mice (Sousa et al., 2009). In these mice, the FRT-flanked stop cassette is removed by Flp-mediated recombination in D1DR+ cells, which leads to GFP expression (Fig 2A). These reporter mice allowed characterization of Flp expression by quantifying GFP+ positive cells and allow comparison of Flp expression with D1DR+ expression using GFP and anti-D1DR staining.

**Figure 2:**
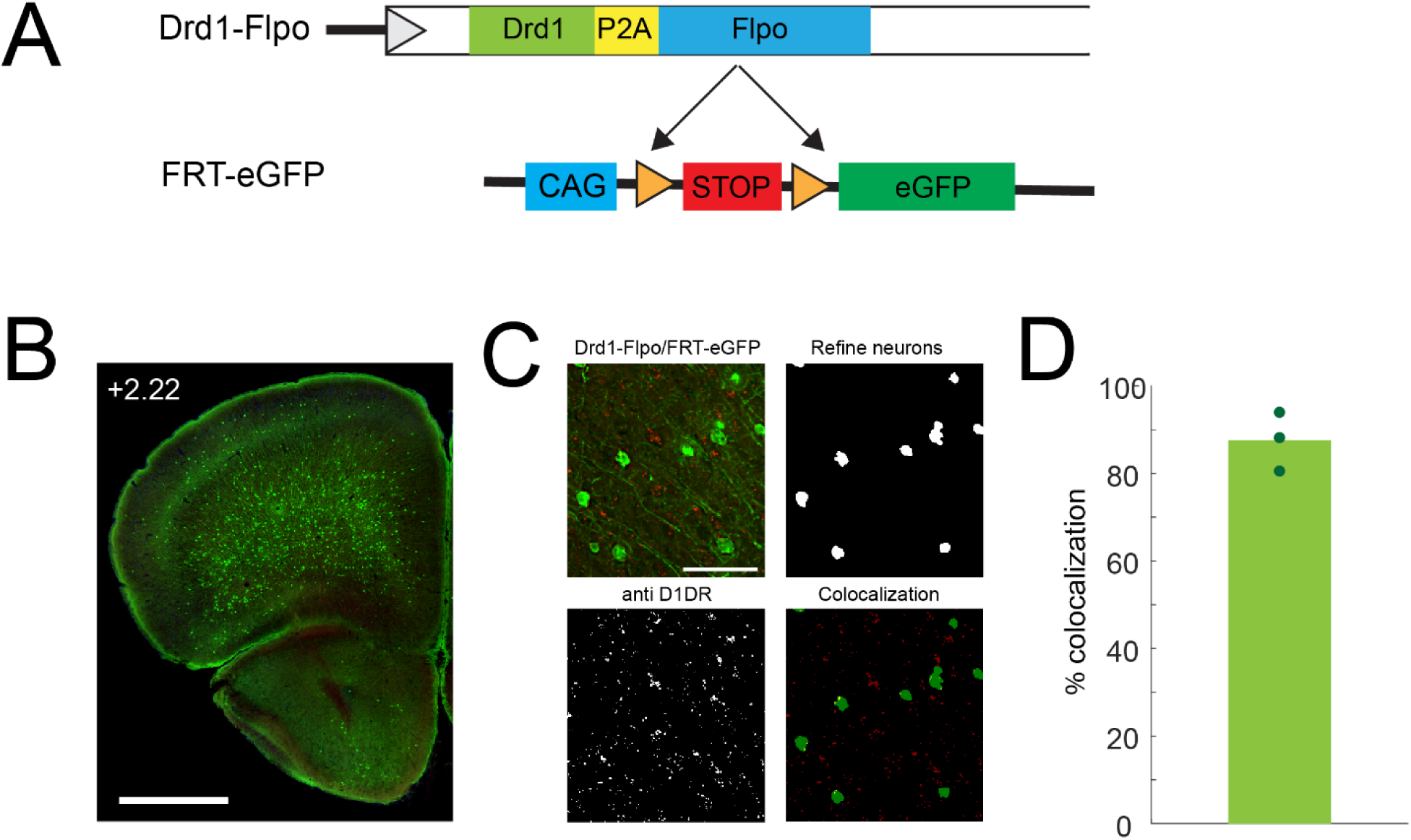
GFP in Drd1-Flpo/FRT-eGFP and anti-D1DR colocalization. (A) Schematic crossing of Drd1-Flpo and Flp-conditional eGFP mouse lines. (B) Five coronal sections of the anterior cortex (AP +1.6 – +0.42) from 3 animals were used to examine colocalization of GFP and anti-D1DR immunoreactivity. Cortical areas of the selected sections were manually defined as regions of interest(scale bar = 1000 um). (C) Colocalization analysis. Imaged with 20x objective (scale bar = 100 um). The cell bodies of GFP expressing neurons were identified with gaussian morphological filtering and active contour (top left). The anti-D1DR immunostaining were converted to adaptive threshold (right bottom). Overlapping pixels were identified to quantify colocalization of signals (top right). (D) Percent colocalization of anti-D1DR and anti-GFP immunostaining in Drd1-Flpo/FRT-GFP cell bodies; each point represents 1 mouse (3 mice).

We found that the Drd1-Flpo/FRT-eGFP mice have strong brain-wide GFP expression in anticipated brain areas such as the frontal cortex (Fig 2B). To quantify Flp expression, we compared GFP+ cells with anti-D1DR+ neurons. Anti-D1DR antibody expression in the striatum was too intense for colocalization; alternatively, because cortical D1DR+ neurons are sparse compared to those in the striatum, it was possible to rigorously colocalize GFP expression with anti-D1DR+ expression (Fig 2C). Strikingly, we found 87.6% ± 3.9 % co-expression of GFP with anti-D1DR positive neurons, indicating specificity of GFP expression (Fig 2D). These results suggest strong Flp colocalization with D1DR+ neurons in the cortex.

### Whole-brain image processing

Next, we examined brain-wide expression of Drd1-Flpo/FRT-eGFP. Qualitative immunostaining analysis on coronally-sectioned (Fig 3) and sagittally-sectioned (Fig 4) brain slices revealed GFP expression in many brain areas. To quantitatively examine brain-wide Flp expression and compare it with D1DR+ expression, we used tissue clearing and light sheet microscopy. We examined whole-brain clearing from Drd1-Flpo/FRT-eGFP in the brains of two mice. Cleared brains were imaged with a light-sheet microscope and then reconstructed as 3D images in MATLAB to generate reconstructions of the coronal or sagittal planes. Serial images were also video-encoded to observe brain-wide expression patterns (Video S1–S2). We mapped atlas-co-registered brains to 1326 sub-regions (Fig 5C). There was no detectable signal of GFP in 438 brain areas. Of the remaining areas, we quantified fluorescence levels of anti-GFP and anti-D1DR in 838 regions. We found a strong positive relatioship (r = 0.53, p < 0.001) between anti-GFP and anti-D1DR intensities within quantified regions (Fig 5D). These analyses suggest that diverse regions of the mouse brain have similar patterns of intensity of Flp-driven eGFP and D1DR immunoreactivity. Taken together, our data support a novel Drd1-Flpo mouse that can be used for circuit-specific investigation of D1DR+ neurons.

**Figure 3:**
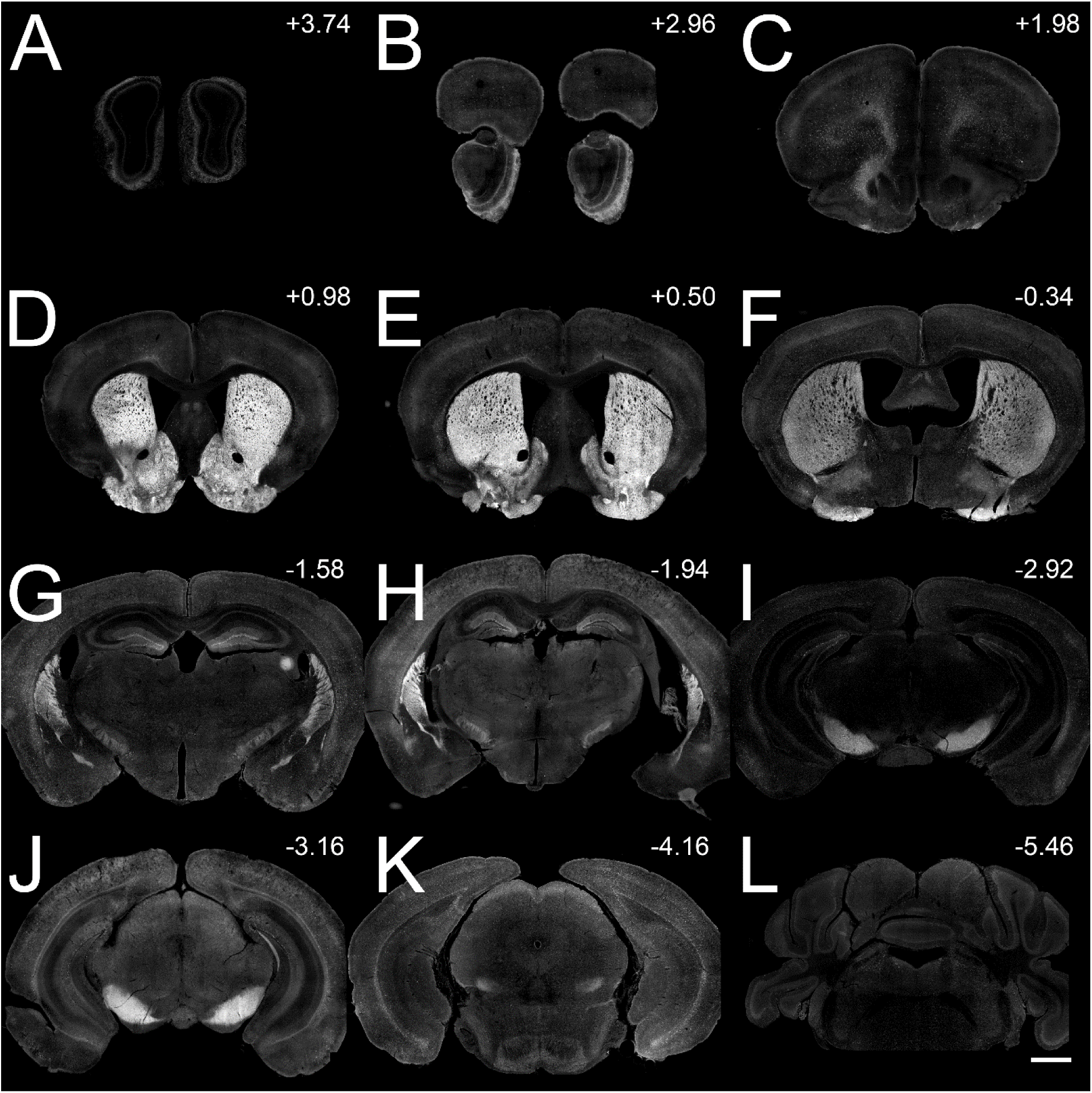
Brain-wide GFP expression. (A–L) GFP expression in coronal sections of Drd1-Flpo/FRT-eGFP mice (3 mice). Anterior to posterior (AP +3.74 to -5.46 mm; scale bar = 1000 um).

**Figure 4:**
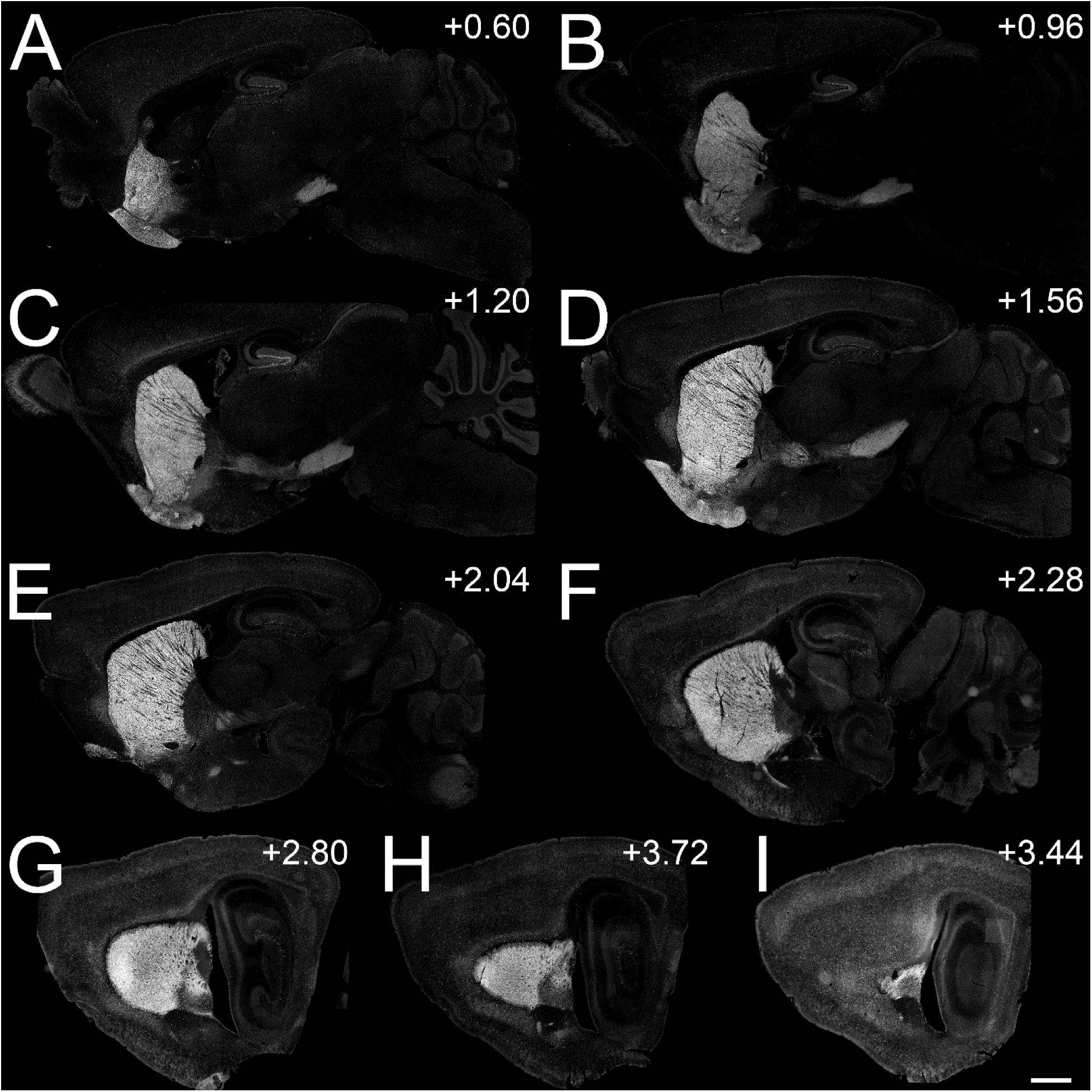
Brain-wide GFP expression. (A–I) GFP expression in sagittal sections of Drd1-Flpo/FRT-eGFP mice (3 mice). Medial to lateral (ML +0.60 to +3.44 mm; scale bar = 1000 um).

**Figure 5:**
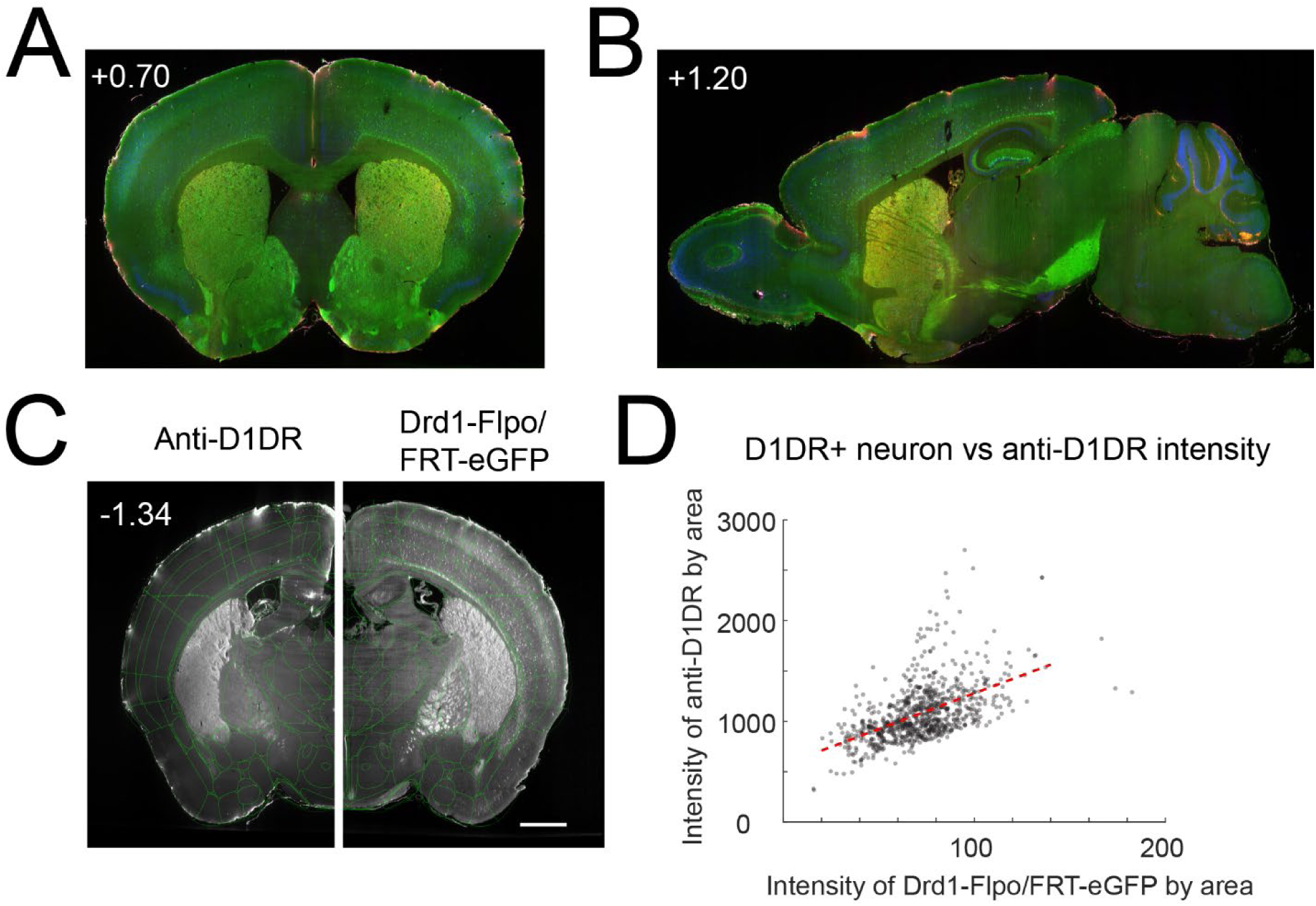
Whole-brain tissue clear imaging. **(A–B)** Examples of reconstructed images from cleared whole-brain imaging in coronal and sagittal planes. **(C)** Example of quantitative analysis from subregions within a coronal 3D image segment mapped to the Allen Brain Atlas (scale bar = 1000 um). **(D)** The relationship between anti-D1DR intensity and anti-GFP intensity in Drd1-Flpo/FRT-eGFP mice per volumetric brain region revealed a strong positive relationship (2 mice).

## Discussion

D1DR+ neurons in the frontal cortex and striatum have been studied in movement, motivation, and cognition; however, their specific roles in these and other behaviors remains unclear. We developed and validated a novel Drd1-Flpo mouse line which facilitates specific interrogation of D1DR+ neurons using conditional and intersectional genetics (Taniguchi et al., 2011; Fenno et al., 2014). We detailed our generation of the Drd1-Flpo line, confirmed Flp recombinase expression and functionality through conditional viral expression of mCherry, and then validated Flp expression and function using a GFP reporter line. Finally, we used whole-brain tissue clearing and light-sheet microscopy to quantify Flp-dependent GFP expression vs. D1DR+ antibodies. The Drd1-Flpo mouse line may be useful for diverse applications specifically interrogating D1DR+ circuits.

D1DR+ neurons are heterogeneous and have functionally- and molecularly-distinct subpopulations (Land et al., 2014; Locke et al., 2018; Anastasiades et al., 2019). Transgenic mice have been a valuable tool for understanding the D1DR+ population in various brain circuits and areas (Lemberger et al., 2007). Previously, our group has used D1-Cre mice to modulate local cortical and striatal networks, which has the potential to improve both cognition and movement (Kim et al., 2017, 2022; Kim and Narayanan, 2019). Although D1DR+ neurons expressing Cre recombinase are a powerful tool, D1-Cre mice are derived from BAC-transgenic lines and thus, can have insertion position effect that can exhibit variability in transgene expression. Furthermore, because D1DR+ neurons contribute to complex brain networks, they can often be expressed alongside several other cell classes, requiring dissection of D1DR+ circuits.

With recent advances in conditional expression and intersectional genetics, multiple genetic switches can be combined to target specific subpopulations of neurons (Fenno et al., 2014). In this study, we generated a new Drd1-Flpo mouse that has functional Flp expression in D1DR+ neurons. The Drd1-Flpo mouse line can be readily combined with other transgenic Cre lines to target more precise neuronal populations, including subpopulations of D1DR+ neurons. Intersectional genetics enables more precise optogenetic manipulation and calcium-sensor-based imaging for more comprehensive analyses of D1DR+ subpopulations. For instance, in the cortex, D1DR+ pyramidal neurons can be studied by crossing Drd1-Flpo mice with pyramidal-neuron-specific Cre driver mice, and in the striatum, Drd1-Flpo mice are expected to facilitate simultaneous imaging of D1DR+ and D2DR+ neurons.

Our report has several limitations. First, we note that D1DRs are expressed on dendrites, axons, and cell bodies of neurons (Levey et al., 1993; Smiley et al., 1994; Bergson et al., 1995; Hersch et al., 1995), and D1DR+ antibody expression can be nonspecific in immunohistochemistry. Since D1DRs are expressed at low levels on the cell body, and their pattern of immunohistochemistry expression may be inconsistent and our colocalization of anti-D1DR and anti-GFP immunostaining was not 100%. Other methods of targeting D1DR+ neurons, such as in situ hybridization, may improve the accuracy of quantifying colocalization; however, this method would also not be capable of identifying D1DR+ axons, fibers, or dendrites. Second, this transgenic mouse line has not been exhaustively behaviorally validated, although mice appear to have normal fertility, growth, and gross behavior. Third, Flp expression may alter D1DR protein levels (Bäckman et al., 2006). In this study we did not examine expression levels of D1DRs. These and general transgenic considerations may be important in designing experiments with Drd1-Flpo mice.

In summary, we report a novel transgenic Drd1-Flpo mouse line. This mouse line can be used to investigate the role of D1DR+ neurons and networks, which could lead to a better understanding and treatment of human diseases affecting movement, motivation, and cognition.

## Code Availability

All code is available at http://narayanan.lab.uiowa.edu/article/datasets.

## Author Contributions

YK and NN designed the experiments. WP designed the mouse. YK and AO performed all experiments and collected data. The data were analyzed by YK and AO. AO, YK, and NN wrote the manuscript.

## Acknowledgements

This work was funded by DOD20037 to Nandakumar Narayanan. Drd1-Flpo transgenic mice were generated at the University of Iowa Genome Editing Facility, which is directed by William Paradee, PhD, and is supported in part by grants from the National Institutes of Health and the Roy J. and Lucille A. Carver College of Medicine.

## Notes

**Declaration of Interests:** The authors declare no competing interests.

### Competing Interest Statement

The authors have declared no competing interest.

